# Polyaxonal A1 amacrine cells form a spatially extended network in primate retina

**DOI:** 10.64898/2026.10.28.762683

**Authors:** Shane Lind Boniec, Fred Rieke, Michael B. Manookin

## Abstract

Inhibitory networks shape information processing in the brain by integrating signals across space, regulating neuronal responses, and controlling the flow of signals through neural circuits. Understanding these computations requires linking the structure of individual inhibitory neuron types to their functional properties and network connectivity. The retina provides a tractable system for meeting this challenge, yet the organization of many retinal inhibitory circuits remains poorly understood. Here, we identify a population of polyaxonal amacrine cells recorded using multielectrode arrays as A1 amacrine cells, a morphologically defined interneuron type in primate retina. This identification is supported by a correspondence between properties of morphologically-identified A1 cells from single cell recordings and putative A1 amacrines in the array recordings. We then identified recursive bistratified ganglion cells in the array recordings based on their electrical coupling to A1 cells, linking this inhibitory network to retinal output pathways. A1 and recursive bistratified cells also share an asymmetry in receptive-field organization, with larger On than Off centers. Finally, A1 spike amplitudes decline during bursts, and analyses of their electrical images suggest that this change may cause some spikes to fail to propagate through their extensive axonal arbors. These findings establish the cellular identity and network organization of a long-range inhibitory circuit in the primate retina and reveal how neuronal architecture, connectivity, and spike dynamics may jointly shape the transmission of visual information.

## INTRODUCTION

Neural computations arise from the interaction of excitation and inhibition in the brain (Isaacson and Scanziani, 2011). Excitatory pathways carry sensory signals, while inhibitory interneurons modulate these signals—sharpening tuning, adjusting gain, and subtracting background inputs (Wehr and Zador, 2003; Isaacson and Scanziani, 2011). In the retina, inhibitory networks perform critical computations beyond simple feedforward relays (Gollisch and Meister, 2010; Diamond, 2017). For example, wide-field amacrine cells convey information from distant regions of the retina, enabling context-dependent suppression and adaptation to overall stimulus statistics (e.g., contrast, motion) (Baccus and Meister, 2002; Ölveczky et al., 2007). By dynamically sculpting the signals that reach ganglion cells, inhibitory circuits help match sensory encoding to the visual environment (Laughlin, 1981). By conveying signals from distant regions into local retinal circuits, they can influence ganglion cell responses to stimuli well beyond the ganglion cells’ receptive-field centers (Ölveczky et al., 2003; Baccus et al., 2008; Cook et al., 1998; Cook and McReynolds, 1998; Hoggarth et al., 2015; Taylor, 1999). In salamander, mouse, and rabbit retina, for example, wide-field amacrine cells transmit signals from the periphery that suppress ganglion cell responses to background motion (Ölveczky et al., 2003; Baccus et al., 2008; Kim and Kerschensteiner, 2017).

How such long-range circuits are organized in the primate retina is poorly understood (Grünert and Martin, 2020). Several types of polyaxonal amacrine cell have been described anatomically, but their functional properties and interactions with other retinal neurons remain largely unknown (Mariani, 1990; Kolb et al., 1992; Dacey, 1989; but see Greschner et al., 2016). A central challenge is linking these known anatomical cell types to their functional roles in visual processing (Dacey et al., 2003; Masland, 2012). Anatomical studies can characterize the morphology of individual cells, whereas large-scale multi-electrode array (MEA) recordings can measure the activity of hundreds of neurons simultaneously and characterize their network interactions but typically can identify them only by physiological properties (Litke et al., 2004; Field and Chichilnisky, 2007). These approaches have therefore provided complementary, but incompletely connected, views of primate retinal circuitry (Li et al., 2015; Rhoades et al., 2019).

One MEA study described a population of polyaxonal amacrine cells with properties consistent with A1 amacrine cells (Greschner et al., 2014), a well-characterized axon-bearing amacrine cell of the primate retina (Dacey, 1989; Stafford and Dacey, 1997; Davenport et al., 2007; Puller et al., 2015; Kim et al., 2022). Here, we provide additional evidence for this identification, including similarities in receptive-field organization, axon arborization patterns, spiking properties, and homotypic electrical coupling. This identification allowed us to find the A1 cell’s ganglion cell partner on the array, the recursive bistratified ganglion cell, based on its electrical coupling to A1 cells and the similarity of its functional properties with anatomically identified recursive bistratified cells (Stafford and Dacey, 1997; Dacey, 2004; Kim et al., 2022; Kling et al., 2024). These cell types both had larger On receptive field centers than Off centers. Finally, A1 spike amplitudes declined during bursts. Simulations based on these amplitude changes suggest that some spikes may fail to propagate through the axonal arbor (Khaliq and Raman, 2005; Monsivais et al., 2005; Debanne et al., 2011; Bucher and Goaillard, 2011). The influence of an A1 cell on downstream neurons therefore depends on both its firing rate and firing pattern (Lisman, 1997). Together, these findings establish the identity and network relationships of A1 amacrine cells and provide evidence that their output may depend not only on firing rate, but also on the temporal pattern of spiking.

## RESULTS

### A1 amacrine cells form a coupled network of axon-bearing interneurons

Anatomical studies have identified ~25 morphological types of amacrine cell in the primate retina (Mariani, 1990; Kolb et al., 1992), and molecular profiling suggests there are more than 30 amacrine types (Peng et al., 2019). However, the physiology of only a few of these cell types is known (Stafford and Dacey, 1997; Greschner et al., 2014; Manookin et al., 2015; Greschner et al., 2016; Grünert and Martin, 2020; Kim et al., 2022). Among these, the A1 amacrine cell is one of the best characterized. A1 cells typically possess multiple axons that branch off the dendritic tree and arborize well beyond it (Dacey, 1989). Whereas their spiny dendrites span approximately 400 µm, their axons extend over several millimeters (Dacey, 1989; Stafford and Dacey, 1997). This distinctive morphology suggests a division of labor: dendrites collect synaptic input and define the receptive field, whereas axons convey spikes over long distances (Davenport et al., 2007).

Greschner et al. (2014) characterized an amacrine population in multi-electrode array (MEA) recordings that shares several properties with the A1 amacrines (described in more detail below). They identified these cells by their radially propagating spikes and showed that their receptive fields are similar in size to their dendritic fields and form an orderly mosaic, and that they have large axonal fields, On–Off light responses, and strong electrical coupling between neighbors. They did not, however, establish a correspondence between these cells and a morphologically identified amacrine cell type. We therefore asked whether the morphologically-identified A1 cells from our patch-clamp recordings share properties with a polyaxonal population identified in array recordings.

We first confirmed key anatomical features in our preparations. Neurobiotin injection into a single A1 cell labeled both the injected cell and a surrounding population of somata (Figure 1A, top), consistent with the tracer coupling reported previously (Stafford and Dacey, 1997). Vertical sections revealed that the dendrites stratified in the middle third of the inner plexiform layer (Figure 1A, bottom), between the two choline acetyltransferase (ChAT)-positive bands that mark the dendrites of starburst amacrine cells (Rodieck and Marshak, 1992).

**Figure 1.**
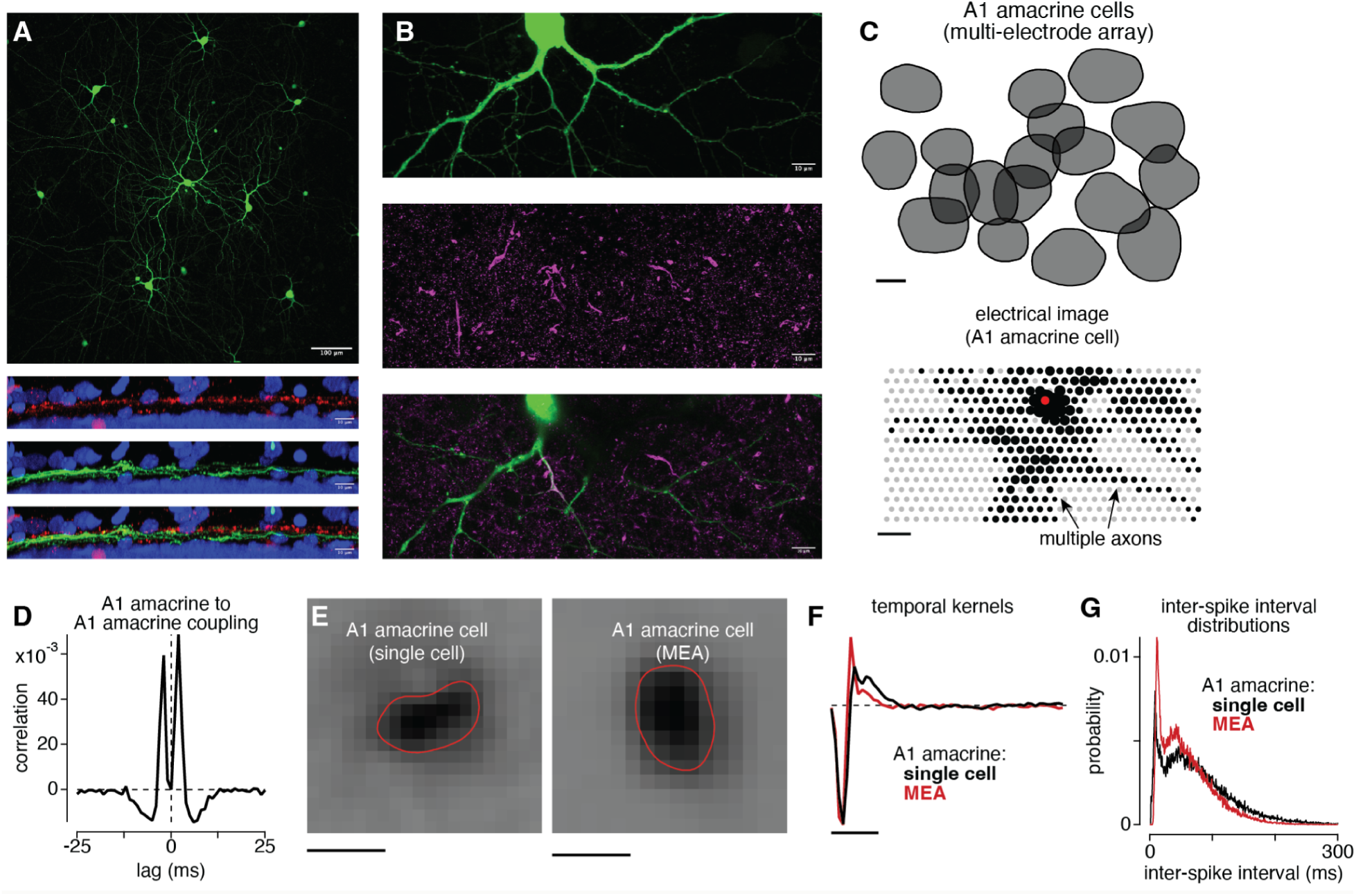
A1 amacrine cells recorded with patch electrodes and on the multi-electrode array. **(A)** Top, maximum-intensity projection of an A1 amacrine cell filled with Neurobiotin. The surrounding labeled somata are tracer-coupled neighbors. Scale bar, 100 µm. Bottom, vertical sections showing A1 dendrites (green) relative to the nuclear layers (DAPI, blue) and the starburst amacrine cell dendrites (ChAT, red). The dendrites stratify in the middle third of the inner plexiform layer. Scale bars, 10 µm. **(B)** Confocal images of a filled A1 amacrine cell (green, top), ankyrin G immunoreactivity marking axon initial segments (magenta, middle), and the merge (bottom). Initial segments lie on processes of the filled cell. The top two images are z-projections and the bottom shows a single frame in the z-plane. Scale bars, 10 µm. **(C)** Top, receptive-field contours for the A1 population recorded on the array in one experiment (18 cells; contours at 0.30 of the peak of the spatial map). Bottom, electrical image of one cell from that population. Gray dots, all electrodes; black markers, electrodes carrying signal, with area proportional to the spike-triggered voltage amplitude; red marker, the electrode eliciting the largest spike amplitude. Arrows mark two of the several axons traversing the array. Scale bars, 200 µm. **(D)** Cross-correlogram of a pair of neighboring A1 cells (1-ms bins; stimulus-driven correlation subtracted, Methods). The two peaks flanking zero lag and the notch at zero are the signature of electrical coupling. **(E)** Spatial receptive fields of an A1 amacrine cell recorded with a patch electrode (left) and of an A1 cell recorded on the array (right), measured with spatiotemporal noise. Red lines show contour fit to the receptive field. Scale bars, 200 µm. **(F)** Temporal kernels of the cells in E. Black, single cell; red, array. Both prefer contrast decrements and are fast and biphasic. Scale bar, 100 ms. **(G)** Probability of spiking as a function of inter-spike interval for the same two cells. Both cells show similar patterns.

To identify potential sites of spike initiation, we immunostained filled cells for ankyrin G, a protein enriched at axon initial segments where it is associated with a high density of voltage-gated sodium channels (Kordeli et al., 1995; Zhou et al., 1998). Ankyrin G immunoreactivity colocalized with processes of the filled cell (Figure 1B). This localization provides structural evidence that the axonal segments may serve as spike-initiation sites in A1 amacrine cells (Zhou et al., 1998).

### A polyaxonal population on the array matches the A1 amacrine cell

We next classified spike-sorted units from multi-electrode array (MEA) recordings using receptive-field mosaics, electrical images, and response kinetics (DeVries and Baylor, 1997; Field and Chichilnisky, 2007; Rhoades et al., 2019). As detailed below, one population shared several features with a polyaxonal amacrine cell population described previously, indicating that the two correspond to the same cell type (Greschner et al., 2014).

First, the identified amacrines had compact receptive fields that tiled the recorded region (Figure 1C, top; 18 cells in the example recording; contour diameter, 391 ± 9 µm). The receptive-field diameter was consistent with the approximately 400-µm dendritic field reported anatomically, as expected if light-driven inputs are collected primarily by the dendrites rather than the extended axonal arbor (Stafford and Dacey, 1997; Davenport et al., 2007; Greschner et al., 2014).

Second, electrical images revealed signals propagating outward from the soma along several distinct paths across the array (Greschner et al., 2014; Li et al., 2015) (Figure 1C, bottom). This spatially extended, propagating activity is consistent with the presence of multiple axons extending beyond the dendritic field. The multiple radiating trajectories distinguished these cells from ganglion cells, which typically have a single axon (Petrusca et al., 2007; Li et al., 2015) (compare Figure 1C and Figure 2A).

**Figure 2.**
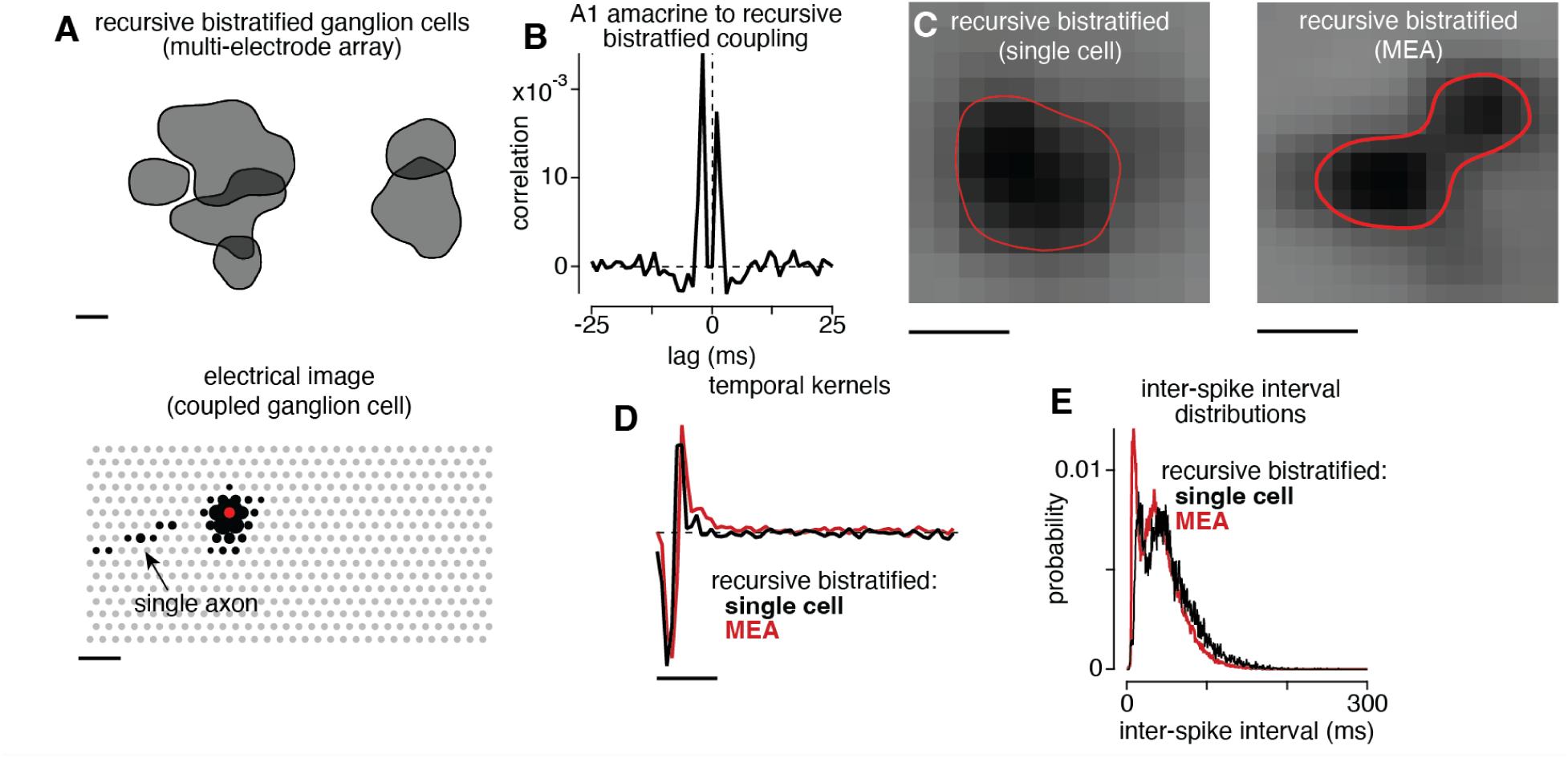
Recursive bistratified ganglion cells on the array and their coupling to A1 cells. **(A)** Top, receptive-field contours for the recursive bistratified population recorded on the array in one experiment (6 cells; contours at 0.30 of the peak of the spatial map). Bottom, electrical image of one cell of that population. Gray dots, all electrodes; black markers, electrodes carrying signal, with area proportional to amplitude; red marker, the soma. The arrow marks the cell’s single axon. Scale bars, 200 µm. **(B)** Cross-correlogram between an A1 amacrine cell and a neighboring recursive bistratified ganglion cell (1-ms bins; stimulus-driven correlation subtracted). Two peaks flank zero lag with a notch between them, indicating electrical coupling between the types. **(C)** Spatial receptive fields of a recursive bistratified cell recorded with a patch electrode (*left*) and on the array (*right*). Red lines show contour fit to the receptive field. Scale bars, 200 µm. **(D)** Temporal kernels of the cells in C. Black, single cell; red, array. Scale bar, 100 ms. **(E)** Inter-spike interval distributions for the same two cells.

Third, the tracer coupling observed anatomically predicts a corresponding signature in the spike trains of neighboring A1 cells. Electrical coupling through gap junctions can produce correlated spiking at short positive and negative time lags, reflecting the propagation of activity between cells in either direction (Brivanlou et al., 1998; DeVries, 1999; Hu and Bloomfield, 2003; Trong and Rieke, 2008). Because transmission is not instantaneous, the resulting cross-correlogram typically exhibits peaks on either side of zero, separated by a central trough. Cross-correlograms from pairs of A1 cells exhibited this characteristic pattern, with two peaks flanking zero at 2 ms and a dip at zero lag (Greschner et al., 2014) (Figure 1D).

This correlation pattern was prominent among nearby cells and declined with distance. Among pairs with receptive-field centers separated by less than 300 µm, nearly all exhibited the double-peaked pattern (44 of 44 and 28 of 30 pairs across two recordings). The fraction decreased to 46%–72% for pairs separated by 300–450 µm and to 4 of 864 (0.5%) for pairs separated by more than 600 µm. Because cells exposed to the same stimulus can exhibit correlated responses even in the absence of direct coupling, we independently estimated stimulus-driven correlations using repeated noise stimuli (Brivanlou et al., 1998; Trong and Rieke, 2008). The double-peaked structure persisted after subtraction of this shared-stimulus component, supporting the interpretation that neighboring A1 cells are electrically coupled (Methods).

The putative A1s in the MEA recordings are distinguished from other recorded amacrine cells based on their rapid and biphasic temporal kernels, Off dominated responses and characteristic inter-spike interval distributions. We tested whether these properties were shared by morphologically-confirmed A1 amacrines in single cell recordings. Spatial receptive fields were similar in size and shape across the two methods (Figure 1E). Temporal kernels were biphasic and rapid, with responses dominated by contrast decrements (Figure 1F). Inter-spike interval distributions also exhibited similar shapes, including a sharp mode at intervals below 25 ms associated with bursts (Figure 1G). The similarity of these response properties provides strong evidence that this population corresponds to A1 amacrine cells.

### A ganglion cell population coupled to A1 cells resembles recursive bistratified cells

Neurobiotin injections into A1 amacrine cells label nearby recursive bistratified ganglion cells, and conversely, injections into recursive bistratified cells label nearby A1 amacrine cells (Stafford and Dacey, 1997; Kim et al., 2022). This suggests that recursive bistratified cells can be identified in MEA recordings based on two-peaked crosscorrelations with A1 amacrines. In this section we further test this identification by comparing functional properties of putative recursive bistratified cells identified in this way with those of morphologically identified cells.

Figure 2 shows six putative recursive bistratified cells identified based on their two-peaked crosscorrelation functions with nearby A1 amacrine cells (Figure 2B). Receptive fields of these cells overlapped minimally, as expected from a single cell type that forms a mosaic. The single axons in their electrical images identify them as retinal ganglion cells.

We further validated the recursive bistratified cell identification by comparing physiological properties measured in array recordings with those measured in single-cell recordings in which cell type was confirmed by recovering morphology after each recording. Spatial receptive fields were similar in size and organization across the two recording methods (Figure 2C), as were the temporal kernels (Figure 2D) and inter-spike interval distributions (Figure 2E). Coupling to putative A1 cells, the sparse mosaic, single-axon electrical images, and agreement with anatomically identified cells together identify this population as recursive bistratified ganglion cells, as in Kling et al. (2024).

### A1 and recursive bistratified cells show On-Off contrast polarity to naturalistic inputs

Receptive fields estimated from spatiotemporal noise are widely used to characterize the visual signals a cell encodes, predict its responses to novel stimuli, and classify cell types in large-scale multi-electrode recordings (DeVries and Baylor, 1997; Chichilnisky, 2001). In particular, the polarity of a cell’s spike-triggered average is commonly used to distinguish On from Off cells (Chichilnisky and Kalmar, 2002; Field and Chichilnisky, 2007). However, this approach is poorly suited to cells that respond to both increments and decrements of light (On-Off cells), for two reasons. First, the spike-triggered average reflects the cell’s dominant contrast polarity as the contribution of spikes evoked by light increments and decrements partially cancel in computing the spike-triggered average (Fairhall et al., 2006; Cantrell et al., 2010). The resulting receptive field emphasizes the polarity that drives more spikes, obscuring contributions from the non-dominant polarity and a key feature that distinguishes On, Off and On-Off cells. In our recordings, both A1 and recursive bistratified cells fired more to light decrements (Off-dominant), so their receptive fields resembled those of Off cells, with the On component largely obscured (Stafford and Dacey, 1997; Davenport et al., 2007; Puller et al., 2015).

A second limitation concerns the static nonlinearity, which describes the relationship between a cell’s mean firing rate and the linear prediction obtained by filtering the stimulus through its receptive field. For an Off-dominant cell, positive predictions correspond to light decrements over the receptive-field center, whereas negative predictions correspond to light increments. A cell that responds to both contrast polarities fires at large predictions of either sign, producing a U-shaped nonlinearity whose two arms reflect the responses to decrements and increments. Conventional white noise, however, rarely generates large linear predictions. Because pixels are randomized independently and updated every 8.3–16.6 ms, neighboring pixels of opposite contrast tend to cancel when filtered by the receptive field. Temporal integration across successive frames further reduces the resulting signal. Consequently, large negative predictions are rare, leaving the negative arm of the nonlinearity—which reflects the On response of an Off-dominant cell—poorly sampled. Responses to the non-dominant polarity may therefore appear weak or absent even in cells that respond to both polarities (Puller et al., 2015).

We used a sparse-noise stimulus to overcome these limitations (Jones and Palmer, 1987). On each frame, a small fraction of pixels (1%) were randomly selected to deviate from the mean gray background; each selected pixel was set to full contrast (+1 for bright, –1 for dark) while all other pixels remained gray. Frames lasted 200 ms, much longer than in the white-noise protocol (8.3–16.6 ms). Under this stimulus, a single dark or bright pixel falling within the receptive field can produce a large filtered contrast of either sign without cancellation by neighboring pixels. The longer frame duration also allows the cell’s temporal response to develop more fully. We computed static nonlinearities from the sparse-noise responses by plotting each cell’s mean firing rate against the filtered stimulus contrast (the projection of the stimulus frame onto the cell’s spatial filter). Both A1 and recursive bistratified cells showed clear On–Off tuning: firing rates increased at both large positive and large negative contrasts (Figure 3A,B, right panels).

**Figure 3.**
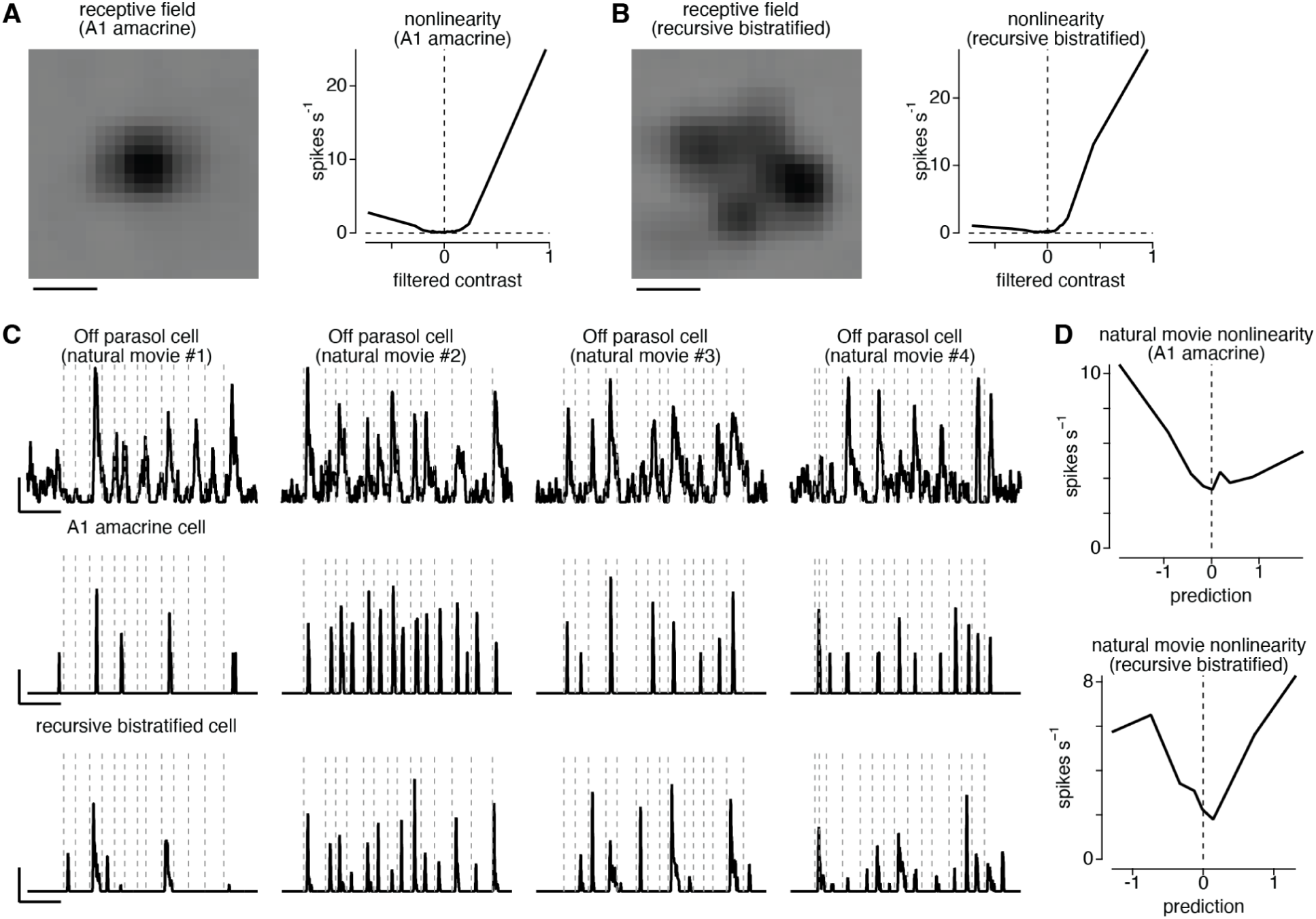
Sparse noise and natural movies reveal On–Off contrast polarity. **(A)** Left, spatial receptive field of an A1 amacrine cell recorded on the array, computed as the spike-triggered average of sparse noise. Right, static nonlinearity for the same cell: mean spike rate as a function of filtered contrast, the projection of each stimulus frame onto the receptive field. Positive filtered contrast corresponds to stimuli that match the receptive field (decrements over the center); negative values correspond to increments. Spike rate rises on both sides of zero. Scale bar, 200 µm. **(B)** As in A, for a recursive bistratified ganglion cell. **(C)** Spike responses to four natural movies (columns) for an Off parasol ganglion cell (top row), an A1 amacrine cell (middle row) and a recursive bistratified ganglion cell (bottom row). Vertical dashed lines mark the timing of saccades. Scale bars: 1 s; 50 spikes s^−1^. **(D)** Static nonlinearities computed from responses to natural movies for an A1 amacrine cell (top) and a recursive bistratified ganglion cell (bottom). The linear prediction was computed by convolving each movie with the cell’s receptive field measured with white noise; spike rate is plotted against this prediction. Spike rate rises on both sides of zero, indicating responses to both contrast polarities.

Thus, despite their Off-dominant spike-triggered averages, both cell types responded to luminance increments and decrements.

We next asked whether this On–Off behavior also appears under naturalistic stimulation. We presented natural movie sequences with simulated eye movements (saccades and fixational drifts) based on human viewing of natural scenes (DOVES database, Van Der Linde et al., 2009). Off parasol ganglion cells responded throughout the movie, with firing rates tracking the changing image (Figure 3C, top row) (Turner and Rieke, 2016; Schottdorf and Lee, 2021). By contrast, A1 and recursive bistratified cells were mostly silent except for brief bursts that often followed saccades (Figure 3C, middle and bottom rows) (compare Greschner et al., 2014; Krishnamoorthy et al., 2017; Krüppel et al., 2023). We computed the nonlinearity for this stimulus by convolving the cell’s spatio-temporal receptive field measured with noise stimuli with the movie sequences, which generated the linear prediction (Chichilnisky, 2001; Karamanlis and Gollisch, 2021). The nonlinearity was then the average spike rate across the range of linear prediction values. Natural images contain large coherent features, so this filtered contrast often takes large positive or negative values. For both A1 and recursive bistratified cells, the resulting nonlinearity was U-shaped (Figure 3D), with higher firing rates at both large positive and large negative contrasts. Thus, the On–Off contrast polarity of these cells, largely hidden in their responses to white noise, emerges clearly under sparse noise and natural movie stimulation.

### On and Off receptive-field centers differ in size in A1 and recursive bistratified cells

Receptive fields of A1 amacrine cells, like many other retinal cells, show an antagonistic center-surround organization (Davenport et al., 2007). To put this receptive field organization in context, we measured responses to spots of increasing diameter and compared A1 cells with three ganglion cell types recorded under the same conditions.

We presented light (increments) and dark spots (decrements) of 18 different diameters, ranging from 18 to 912 µm, centered on the receptive field (contrast, ±0.5; duration, 0.5 s). A1 cells responded to both the contrast increments and decrements (Figure 4A). Both responses were transient. The decrement response increased with spot diameter up to approximately 250 µm, then declined by roughly half for the largest spots (Figure 4B, filled circles). Thus, extending the stimulus beyond the diameter that elicited the maximal response reduced rather than further increased the response, demonstrating substantial surround suppression.

**Figure 4.**
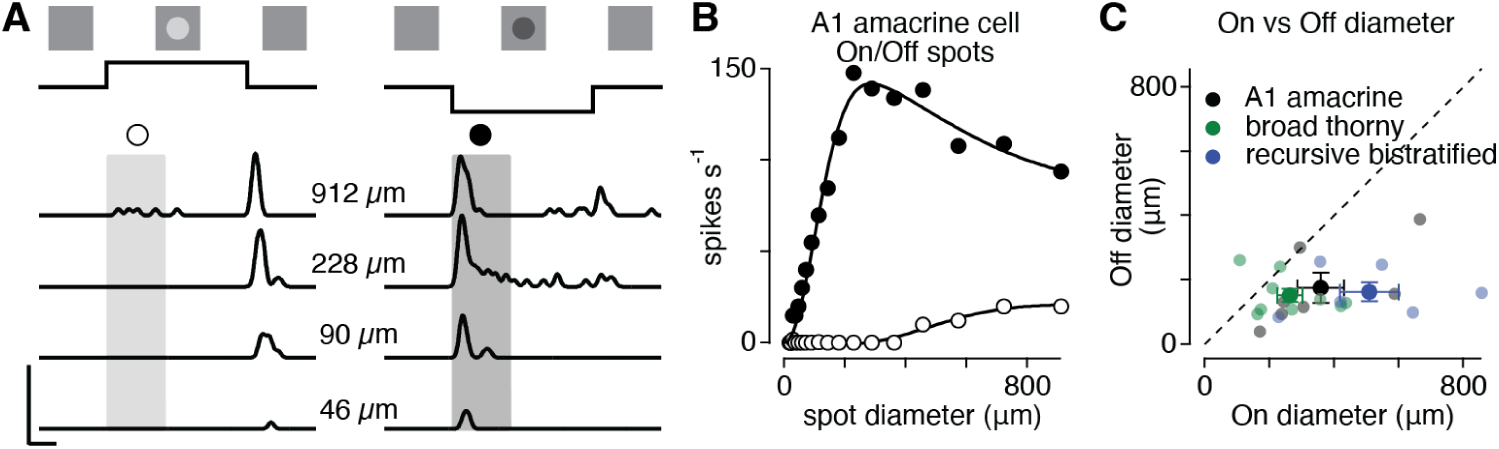
Spatial receptive-field structure measured with spots of varying diameter. **(A)** Spike responses of an A1 amacrine cell to light (left) and dark (right) spots (contrast, ±0.5; 0.5-s duration) of four diameters, centered on the receptive field. The schematic above the traces shows the stimulus time course. The open circle marks the contrast increment and the filled circle marks the decrement. Shading marks the windows over which spike rates were averaged. Scale bars, 500 spikes s^−1^ and 100 ms. **(B)** Average spike rate against spot diameter for the cell in A, across 18 diameters (18–912 µm). Filled circles, response to the onset of dark spots (Off); open circles, response to the onset of bright spots of the same diameters (On). Curves are fits of a difference-of-Gaussians area-summation model. **(C)** Receptive-field center diameter estimated from the On response against the estimate from the Off response, for each A1 amacrine (black), broad thorny (green) and recursive bistratified cell (blue). Center diameter is the spot diameter at which the area-summation curve first reached 1 − 1/e of its maximum. Large symbols with error bars, mean ± SEM per type; dashed line, unity. On centers exceeded Off centers in A1 (p = 0.031, n = 7) and recursive bistratified cells (p = 0.031, n = 6) but not in broad thorny cells (p = 0.055, n = 9; Wilcoxon signed-rank tests).

Although A1 cells respond to both contrast increments and decrements, these responses need not originate from spatially equivalent regions of the receptive field. Indeed, in the example shown in Figure 4B, the increment response peaked at a substantially larger spot diameter than the decrement response (approximately 910 versus 230 µm). This difference suggests that the spatial extent of the receptive-field center depends on contrast polarity (Chichilnisky and Kalmar, 2002; Ravi et al., 2018). We therefore quantified the On and Off center sizes separately in each On–Off cell and compared the resulting estimates within individual cells (Figure 4C).

We defined the center diameter as the spot diameter at which the area-summation curve first reached (1-1/e) of its maximum response. Applying this criterion separately to the On and Off responses allowed us to compare their spatial scales within each cell, independently of differences in absolute response magnitude. We measured the On response at the onset of bright spots and the Off response at the onset of dark spots, so that both started from the mean background. The On receptive-field center in A1 cells was larger than the Off center (358 ± 72 versus 175 ± 47 µm; On larger in 6 of 7 cells; Wilcoxon signed-rank test, p = 0.031). The same asymmetry was present in recursive bistratified ganglion cells (509 ± 91 versus 163 ± 30 µm; 6 of 6 cells; p = 0.031).

Thus, A1 and recursive bistratified cells share an asymmetry in the spatial extent of their On and Off responses. Because these comparisons were performed within individual cells, the observed differences cannot be attributed simply to variation in absolute receptive-field size across the sampled populations. The shared asymmetry in the two coupled cell types raises the possibility that they share aspects of the circuitry or synaptic mechanisms governing the spatial integration of On and Off signals (DeVries, 1999; Cooler and Schwartz, 2021).

### A chirp stimulus distinguishes the four cell types

The four cell types described above exhibited similar spatial receptive fields. To compare their temporal response properties and contrast coding we used a chirp stimulus that combines luminance steps, a temporal-frequency sweep, and a contrast sweep. This provides a compact assay of response polarity, temporal tuning, and contrast sensitivity. This stimulus has been used to distinguish retinal ganglion cell types in mouse retina (Baden et al., 2016).

The stimulus began with full-field light and dark steps, followed by a 10-s sinusoidal frequency sweep from 0 to 10 Hz at full contrast and an 8-s contrast sweep from 2% to 100% contrast at 2 Hz (Figure 5A). Differences in the responses of the four cell types to this stimulus provided complementary physiological signatures with which to distinguish the populations.

**Figure 5.**
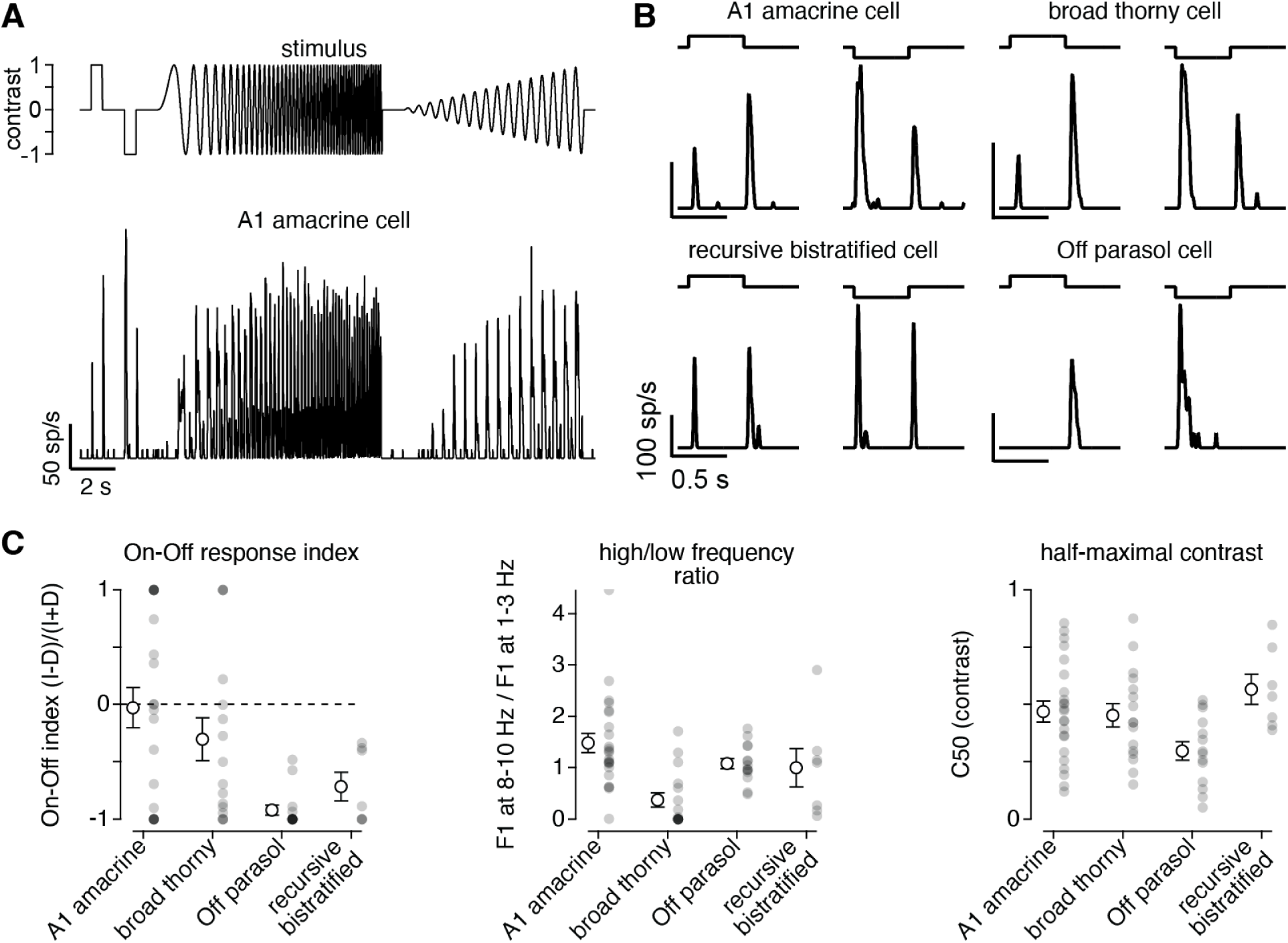
Temporal and contrast coding measured with a chirp stimulus. **(A)** Top, the stimulus: 500-ms light and dark steps at full contrast, a 10-s linear frequency sweep from 0 to 10 Hz at full contrast, and an 8-s contrast sweep at 2 Hz from 2% to 100%. Background, half-maximal intensity. Bottom, the response of an A1 amacrine cell to the full stimulus. Scale bars, 50 spikes s^−1^ and 2 s. **(B)** The step portion expanded, for one cell of each type. The stimulus time course is drawn above each trace. A1, recursive bistratified and broad thorny cells respond at the onset and the offset of both steps; Off parasol cells respond to contrast decrements only. **(C)** One summary value per cell, by type. Left, On-Off index, (I − D)/(I + D), where I is the response to the light-step onset and D the response to the dark-step onset; −1, dimming only; +1, brightening only. Middle, ratio of the F1 response amplitude at 8–10 Hz to that at 1–3 Hz during the frequency sweep. Right, half-maximal contrast (C_50_) of the contrast sweep. Gray circles, individual cells; open circles with error bars, mean ± SEM. A1, n = 23 (22 for the On-Off index, which is undefined for a cell whose rate rose after neither onset); broad thorny, n = 16; Off parasol, n = 14; recursive bistratified, n = 7. Each measure differed across types (Kruskal–Wallis; left, p < 0.001; middle, p < 0.001; right, p = 0.026); comparisons with Off parasol cells are given in the Results.

Responses to the initial luminance steps revealed clear differences in the relative strength of On and Off responses (Figure 5B). A1, recursive bistratified, and broad thorny cells responded at both the onset and offset of the light step and at both transitions of the dark step, indicating sensitivity to both contrast increments and decrements. In contrast, Off parasol cells responded predominantly at light offset and dark-step onset, with little activity at the opposite transitions, consistent with their predominantly Off response polarity (Chichilnisky and Kalmar, 2002). To compare polarity across cells, we computed an On-Off index (Carcieri et al., 2003), (I − D)/(I + D), where I is the response to the onset of the light step and D the response to the onset of the dark step. Both onsets start from the mean background, so the index is not affected by adaptation during the steps. The index is −1 for a cell that responds only to dimming and +1 for one that responds only to brightening (Methods). Off parasol cells had an index near −1 (−0.96 ± 0.03, n = 14; Figure 5C, left). The three On–Off types lay between the extremes, and the index differed across types (Kruskal–Wallis, p < 0.001).

To compare temporal tuning across cells, we took the ratio of the response amplitude for stimulus frequencies (F1) between 8 and 10 Hz to that between 1 and 3 Hz, within the 0–10 Hz range swept in every recording (Methods). A ratio above 1 indicates a cell that follows fast modulation better than slow modulation; a ratio below 1 indicates low-pass tuning. The ratio differed across types (Kruskal–Wallis, p < 0.001; Figure 5C, middle). Broad thorny cells were strongly low-pass (0.38 ± 0.14, n = 16; rank-sum against Off parasol cells, p = 0.002). In contrast, A1 (1.48 ± 0.19, n = 23), recursive bistratified (1.00 ± 0.37, n = 7) and Off parasol cells (1.07 ± 0.10, n = 14) responded at least as strongly at high as at low frequencies, and neither A1 nor recursive bistratified cells differed from Off parasol cells (p = 0.10 and p = 0.69). These response profiles indicate that A1, recursive bistratified, and Off parasol cells sustain or increase their responses across a relatively broad range of temporal frequencies (for parasol cells, see also Lee et al., 1994; Benardete and Kaplan, 1999), whereas broad thorny cells preferentially respond at lower frequencies.

The contrast sweep revealed additional differences in how the four cell types encode stimulus strength. We summarized each contrast-response function by its half-maximal contrast (C_50_), the contrast at which the response first reached half its maximum, measured without fitting a model (Methods). C_50_ differed across types, and this difference came from Off parasol cells (Kruskal–Wallis, p = 0.026; Figure 5C, right). Off parasol cells reached half-maximal response at the lowest contrast (0.30 ± 0.04, n = 14), consistent with a high contrast gain (Kaplan and Shapley, 1986; Croner and Kaplan, 1995). A1, broad thorny, and recursive bistratified cells all showed significantly higher C_50_ values, consistent with a lower contrast gain compared to parasol cells (A1: 0.47 ± 0.04, n = 23, p = 0.025; broad thorny: 0.45 ± 0.05, n = 16, p = 0.048; recursive bistratified: 0.57 ± 0.07, n = 7, p = 0.003; rank-sum tests against Off parasol cells), and these cells did not differ from one another (Kruskal–Wallis, p = 0.47). These lower contrast gain values are consistent with their lower maintained firing rates (pooled median 0 vs. 8.9 spikes/s for Off parasol cells; p = 0.002). With little ongoing activity, weak contrasts would leave these cells below spike threshold, whereas the maintained firing of Off parasol cells allows smaller modulations to change their output.

### A1 and recursive bistratified cells are not robustly direction selective

A recent study reported direction selectivity in both A1 amacrine cells and recursive bistratified ganglion cells (Kim et al., 2022). In particular, recursive bistratified cells exhibited a mean direction-selectivity index of 0.82 in response to moving bars, consistent with strong directional tuning. This finding has important implications for the functional organization of the network described above: if both electrically coupled cell types exhibit strong direction selectivity, their interactions could contribute to a directionally tuned circuit. We therefore tested direction selectivity in both cell types using classical motion stimuli (Barlow and Levick, 1965; Oyster and Barlow, 1967; Vaney et al., 2012) and two recording configurations. Broad thorny and Off parasol ganglion cells, which are not generally considered direction selective, served as controls for the degree of apparent tuning measured under the same conditions (Puller et al., 2015; Manookin et al., 2018).

We presented drifting gratings and moving bars in 8–12 directions and measured responses in A1 amacrine, recursive bistratified, broad thorny, and Off parasol cells. These stimuli allowed us to assess directional tuning using two distinct forms of visual motion. Drifting gratings probe responses to spatially periodic motion, whereas moving bars provide a spatially localized stimulus with distinct leading and trailing edges. We quantified directional tuning using both the direction-selectivity index (DSI) and orientation-selectivity index (OSI), allowing us to distinguish a preference for motion in one direction from a preference for a particular orientation regardless of motion direction.

A1 cells responded robustly to drifting gratings moving in all tested directions, with little apparent dependence on direction (Figure 6A). To quantify direction selectivity, we calculated the vector sum of responses across stimulus directions, with each response contributing a vector oriented along its corresponding direction of motion and weighted by response magnitude (see Methods). The direction of the resultant vector indicates the preferred direction, whereas its magnitude, normalized by the mean response across directions, provides the direction-selectivity index (DSI) (Taylor and Vaney, 2002). Thus, a DSI near zero indicates little directional bias, whereas larger values indicate stronger direction selectivity. DSI values were small across A1 cells, indicating at most weak directional tuning. Moving bars produced a similar result (Figure 6B). A1 cells responded to both the leading and trailing edges of the bar across the tested directions, with no consistent preference for a particular direction of motion.

**Figure 6.**
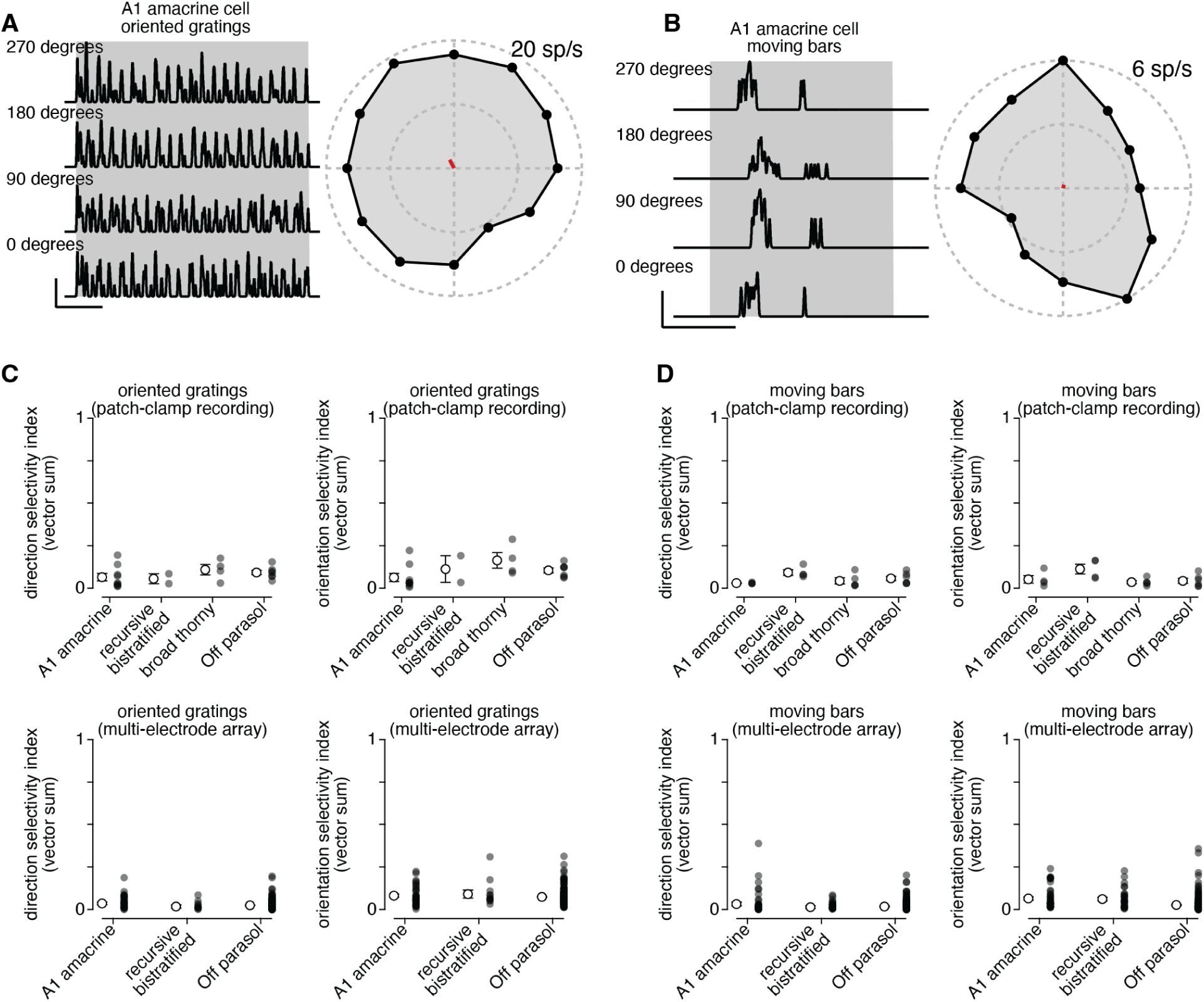
Direction and orientation selectivity across four cell types. **(A)** Left, firing rate of an A1 amacrine cell to gratings drifting in 12 directions; four directions are shown. Shading marks the period of grating motion. Right, mean firing rate against direction for the same cell. The red vector from the origin gives the direction and magnitude of the vector-sum preference; the outer dashed circle is 20 spikes s^−1^ (scale bars: x, 1 s; y, 50 spikes s^−1^). **(B)** As in A, for bars moving in 12 directions. The outer dashed circle is 6 spikes s^−1^ (scale bars: x, 1 s; y, 50 spikes s^−1^). **(C)** Direction-selectivity index (left) and orientation-selectivity index (right) measured with drifting gratings, from patch-clamp recordings of four types (top) and array recordings of three types (bottom). Small symbols, individual cells; open circles with error bars, mean ± SEM. Patch-clamp: A1, n = 9; recursive bistratified, n = 3; broad thorny, n = 4; Off parasol, n = 7. Array: A1, 34 cells across 2 experiments; recursive bistratified, 12 cells across 2 experiments; Off parasol, 167 cells across 2 experiments. In patch-clamp recordings, neither A1 nor recursive bistratified cells differed significantly from Off parasol cells in DSI, for drifting gratings (A1, p = 0.21; recursive bistratified, p = 0.33; Wilcoxon rank-sum test). The same held for OSI (p = 0.055 and p = 1.00). On the array, neither type differed from Off parasol cells in DSI (p = 0.079 and p = 0.49) or OSI (p = 0.68 and p = 0.55). Patch-clamp tests use the measured index; array tests use the bias-corrected index, while the plotted values are raw in both (Methods). Neither index differed across types in the patch-clamp data (Kruskal–Wallis, p = 0.32 and p = 0.097) or in the array data (p = 0.15 and p = 0.78). **(D)** As in C, for the moving-bar data. Patch-clamp: A1, n = 4; recursive bistratified, n = 4; broad thorny, n = 5; Off parasol, n = 6. Array: A1, 47 cells across 3 experiments; recursive bistratified, 29 cells across 3 experiments; Off parasol, 176 cells across 3 experiments. In patch-clamp recordings, DSI did not differ significantly for A1 or recursive bistratified cells relative to the Off parasol controls (p = 0.067 and p = 0.35; Wilcoxon rank-sum test). This was also true for OSI (p = 0.76 and p = 0.067). On the array, DSI again did not differ (p = 0.91 and p = 0.90), but A1 and recursive bistratified cells had higher OSIs than Off parasol cells (p < 0.001 and p < 0.001); because a moving bar has leading and trailing edges, this reflects a preferred axis of motion rather than a preferred direction. Neither index differed across types in the patch-clamp data (p = 0.076 and p = 0.13). In the array data DSI did not (p = 0.99) but OSI did (p < 0.001; Results).

Population analyses confirmed these observations. In patch-clamp recordings, mean DSI values for A1 and recursive bistratified cells to both gratings and bars were small and comparable to those measured in the untuned control population. These populations did not differ significantly in DSI (Kruskal–Wallis test, p = 0.32 for gratings and p = 0.076 for bars) or OSI (p = 0.097 for gratings and p = 0.13 for bars).

MEA recordings provided a larger sample with which to test these findings. Across A1, recursive bistratified, and Off parasol cells, direction-selectivity indices (DSIs) were generally low, with substantial overlap among populations (Figure 6C,D). DSI values below approximately 0.2 fell within the range observed in the untuned Off parasol controls and therefore did not provide compelling evidence of direction selectivity (see Mazurek et al., 2014). In summary, under the conditions of our experiments neither A1 amacrine nor recursive bistratified ganglion cells showed robust direction or orientation selectivity (see Discussion).

### Spike amplitude adaptation affects signal propagation in A1 axonal arbors

Thus far, we have considered all of the spikes recorded in A1 cells as equivalent events. In practice, however, our recordings indicate that spike amplitude can vary substantially within bursts of spikes (Fee et al., 1996; Quirk and Wilson, 1999) (Figure 7A). For example, during high-frequency firing, the second and third spikes in an A1 burst often have smaller amplitudes than the first (Figure 7A,B). This amplitude adaptation was also apparent in responses to spatiotemporal noise, and it was markedly larger in A1 cells than in Off parasol cells under the same stimulus conditions (Figure 7C). These observations raised the possibility that the smaller spikes might propagate less efficiently along the cell’s long axonal arbor (Grossman et al., 1979; Debanne et al., 1997; Khaliq and Raman, 2005; Monsivais et al., 2005; Debanne et al., 2011)? In other words, could the spike train arriving at distant targets differ from the train recorded near the soma when bursts occur? To investigate this, we examined the spatial extent of each spike’s extracellular signature across the multi-electrode array.

**Figure 7.**
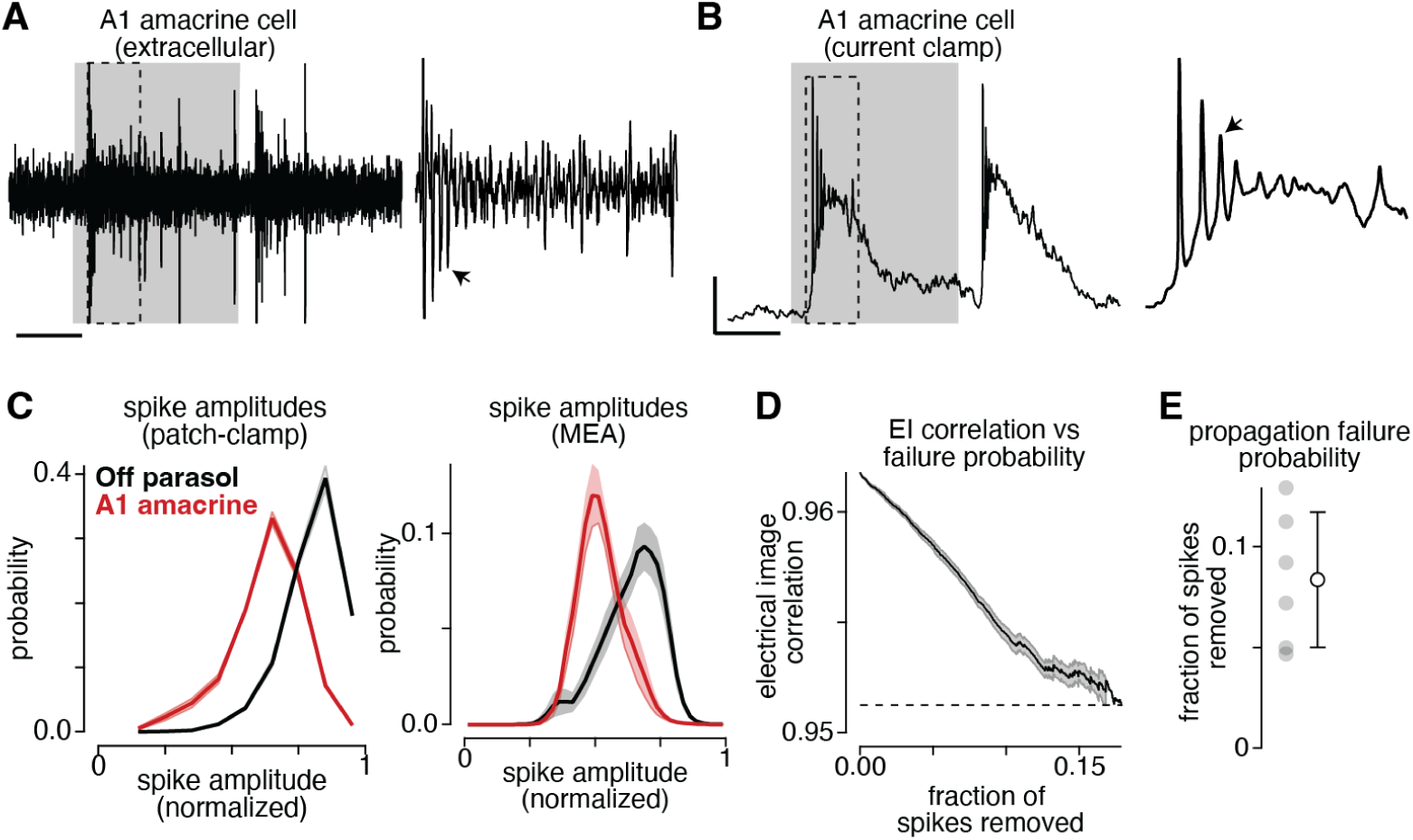
Spike amplitude adaptation within a burst, and its consequences for axonal propagation. (A,B) Extracellular (A) and whole-cell current-clamp recordings (B) showing the responses of an A1 amacrine cell to a light step. The timing of the light step is indicated by the gray region. Spike height varied considerably during the step. Scale bars: x-axis, 100 ms; y-axis, 10 mV. **(C)** Distributions of spike amplitude for Off parasol (*black*) and A1 amacrine cells (*red*). Spikes were normalized to the largest spike amplitude within each epoch. Data are shown for patch-clamp (*left*) and array recordings (*right*). **(D)** Converting the difference between electrical images into a number of propagation failures, for one example cell. The curve is the correlation between the cell’s full electrical image and the image built from its first burst spikes, as simulated failures are introduced one spike at a time (mean of 20 simulations; shading, ± SEM). Images were compared by Pearson correlation and zero-centered by subtracting, for each spike, the mean array signal over a 3-ms (60-sample) segment ending 4 ms (80 samples) before the spike. A failure was simulated by replacing the array data during that spike with the data recorded 4 ms earlier. Dashed line indicates the correlation value measured for the third spike across bursts. **(E)** Fraction of spikes that must fail for the first-spike image to match the third-spike image, for six individual A1 amacrine cells (*left*) and the mean ± SEM (*right*).

For each A1 cell, we computed separate electrical images for spikes at different positions within a burst (e.g., the first and third spikes) and compared each image with the cell’s full electrical image, calculated by averaging across all spikes (Litke et al., 2004; Li et al., 2015). We quantified similarity using the Pearson correlation coefficient, which measures the correspondence between the spatial patterns of extracellular voltage signals across electrodes. Across cells, the electrical image for the third spike in a burst showed a lower correlation with the full electrical image than was the image for the first spike (r_1st_ = 0.962 ± 0.005, r_3rd_ = 0.951 ± 0.006; mean ± SEM across n = 20 repeats; Figure 7D). This difference was consistent across all cells: third-spike images were less correlated with the full image than first-spike images (Wilcoxon signed rank test, p = 0.016). Thus, later spikes were associated with a measurably different spatial pattern of extracellular signals across the array.

To determine how many spikes would need to fail to propagate to account for this difference, we performed a simulated spike-removal analysis. Starting from the first-spike electrical image, we randomly removed spikes by replacing their multi-electrode waveform with baseline noise from 4 ms earlier. Each replacement mimics the loss of a spike’s signal on the array while preserving the recording noise. After each removal, we recomputed the image’s correlation with the full image. As more spikes were removed, the correlation declined monotonically (Figure 7D): with no removals, the correlation was ~0.962, and after removal of 16% of the spikes it fell to ~0.951 - matching the measured third-spike correlations and defining an “equivalent removal fraction.” Across A1 cells, this equivalent removal fraction ranged from 0.047 to 0.129, with a mean of ~0.084 (Figure 7E). In other words, the third-spike image in each cell looked like the first-spike image with roughly 8% of its spikes missing.

In summary, the electrical-image signatures of later A1 burst spikes are measurably smaller than those of early spikes. While this does not prove that individual spikes fail to reach distal axonal branches, it suggests that A1 axonal output may not be a faithful copy of somatic firing. If later spikes in a burst are indeed lost or attenuated along the arbor, then somatic firing rate alone overestimates the signals delivered to targets (Khaliq and Raman, 2005; Monsivais et al., 2005). In that case, the temporal pattern of A1 cell firing – not just the overall spike count – would determine its influence on downstream neurons (Lisman, 1997).

## DISCUSSION

We set out to link a morphologically defined retinal interneuron to a population recorded using a multi-electrode array (MEA). Four independent lines of evidence identify this population as the A1 amacrine cell. First, its receptive fields form a mosaic with a characteristic diameter consistent with the anatomical extent of A1 dendrites (Figure 1). Second, its electrical images reveal multiple axonal trajectories extending millimeters beyond the receptive-field center, a distinctive feature of this polyaxonal cell (Litke et al., 2004; Greschner et al., 2014). Third, neighboring cells exhibit short-lag correlation patterns consistent with the electrical coupling predicted by tracer-injection studies (Stafford and Dacey, 1997; Puller et al., 2020; Kim et al., 2022). Fourth, their receptive fields, temporal kernels, and interspike-interval distributions match those of anatomically identified A1 cells recorded using patch clamp (Figure 1). These features also match the polyaxonal population characterized in earlier MEA recordings, for which the morphological identity had not been established (Greschner et al., 2014).

Having established the identity of the A1 population, we could use its coupling relationships to identify its ganglion-cell partner. The coupled population matched the properties of anatomically identified recursive bistratified ganglion cells (Figure 2; Stafford and Dacey, 1997; Kim et al., 2022; Kling et al., 2024). This distinction is important because the coupling relationship provides evidence for the identity of the partner population only after the A1 cells have been independently identified; it does not, by itself, establish the identity of either population. Our identification of recursive bistratified cells is therefore grounded in comparisons with anatomically identified cells, while the MEA recordings extend these observations to larger populations. Together, these findings establish a physiological framework for identifying A1 amacrine cells and their coupled partners in large-scale recordings.

### Polarity-dependent spatial integration in natural scenes

A1 amacrine and recursive bistratified ganglion cells integrate brightening over a wider region than darkening: within individual cells, the On center was larger than the Off center, by roughly twofold (Figure 4B). Broad thorny cells, another On–Off type, showed no such difference. The asymmetry runs in the same direction as that between the major primate pathways, in which On parasol and midget cells have larger receptive fields than their Off counterparts (Dacey and Petersen, 1992; Chichilnisky and Kalmar, 2002). That difference has been interpreted as an adaptation to natural scenes, which contain more dark than light contrast at the spatial scales sampled by ganglion cells; sampling dark features more finely than light ones matches the retina’s resources to that excess of darkness (Field, 1987; Dong and Atick, 1995; Ratliff et al., 2010; Gjorgjieva et al., 2014). In A1 and recursive bistratified cells, the same trade-off appears within single cells rather than between two cell types.

Because the On and Off centers differ in extent, the two polarities should be selective for features of different sizes. Off responses would be strongest for dark features that roughly fill the narrow Off center, whereas On responses would grow with bright regions up to about twice that width. A natural image patch and its contrast-reversed counterpart would therefore not be encoded symmetrically: the same spatial arrangement of light and dark would be read at a fine scale where it is dark and at a coarse scale where it is bright. Given the excess of dark contrast in natural scenes, this arrangement would give dark structure high spatial resolution while pooling sparser bright structure over a larger area. Which features dominate the response to a given scene would then depend on how dark and bright structures are distributed across spatial scales in that scene. For A1 cells, whose axons distribute their output over millimeters, this size selectivity would carry over to the long-range signal they send to distant circuits.

These predictions rest on center sizes measured with uniform spots, which describe the area over which each polarity is integrated but not how signals are combined within that area. Receptive fields that pool bipolar input are built from smaller subunits whose rectified outputs make responses sensitive to spatial structure that a uniform spot does not probe (Demb et al., 2001; Hochstein and Shapley, 1976; Schwartz et al., 2012; Freeman et al., 2015). Natural images contain substantial structure within a single receptive field, and nonlinear integration across subunits strongly shapes responses to them, in ways that can differ between On and Off pathways (Turner and Rieke, 2016). A wide On center built from small, strongly rectified subunits would remain sensitive to fine bright features within it, whereas one built from larger or more linear subunits would average them away. The sizes and nonlinearities of the On and Off subunits, not center size alone, will therefore determine how the asymmetry shapes responses in natural vision. Mapping these subunits directly, or comparing responses to natural image patches with responses to linear-equivalent stimuli (Turner and Rieke, 2016; Turner et al., 2018), is a necessary next step, together with models that allow both the spatial extent and the subunit structure of integration to differ between polarities.

Several mechanisms could produce the asymmetry. On and Off bipolar inputs may differ in spatial extent, polarity-specific inhibition may shape integration differently for the two signs (Manookin et al., 2008; Werblin, 2010), or electrical coupling between A1 and recursive bistratified cells may broaden the effective receptive field more for one polarity than the other (DeVries, 1999; Cooler and Schwartz, 2021). The shared asymmetry in two coupled cell types is consistent with a contribution from coupling but does not establish it. If coupling contributes, the difference between On and Off center sizes should covary with a cell’s coupling to its neighbors, a relationship that could be examined with the existing pairwise coupling measurements, provided coupling strength and receptive-field asymmetry can be estimated reliably for the same cells. A positive relationship would support a network contribution, although it would not exclude differences in bipolar input or inhibition.

### Direction selectivity

A previous study reported strong direction selectivity in both A1 amacrine and recursive bistratified ganglion cells (Kim et al., 2022). We did not see strong direction selectivity in these cells with either drifting gratings or moving bars, using either patch-clamp or MEA recordings (Figure 6). In both populations, directional tuning was weak and comparable to that of the untuned controls: Off parasol cells in both recording configurations and broad thorny cells in patch-clamp recordings. While the origin of this discrepancy remains unresolved, it suggests that any direction selectivity of these primate cell types is less robust to recording conditions than that observed in direction-selective ganglion cells of rabbit and mouse retina, which remain selective across a wide range of stimulus speeds, contrasts, sizes and light levels, and in both single-cell and multi-electrode recordings (Barlow and Levick, 1965; Oyster, 1968; Wyatt and Daw, 1975; Grzywacz et al., 1994; Taylor and Vaney, 2002; Weng et al., 2005; Elstrott et al., 2008; Lipin et al., 2015; Poleg-Polsky and Diamond, 2016; Sabbah et al., 2017; Yao et al., 2018; Wei, 2018).

### Spike propagation efficiency

A1 cells exhibit pronounced spike-amplitude adaptation within bursts, and the electrical images associated with the first and third spikes differ in their similarity to the full electrical image (Figure 7). These findings establish that the spatial pattern of extracellular signals associated with a spike depends on its position within a burst, but they do not establish the underlying mechanism. The reduced correlation is consistent with approximately 8% of later spikes failing to propagate, but uniform amplitude scaling could produce a similar effect without any spikes being lost along the axonal arbor. Distinguishing these explanations requires determining whether signal loss increases with distance along the axon and establishing that the analysis does not produce a comparable apparent failure fraction in a control population without amplitude adaptation.

If subsequent analyses support genuine propagation failures, this would suggest that A1 axons do more than faithfully distribute the spike output generated near the soma. The conventional view distinguishes the roles of dendrites, which collect synaptic input, and axons, which distribute the resulting spike output (Davenport et al., 2007). However, if spikes occurring later in a burst are less likely to reach distant axonal locations, a burst recorded as three somatic spikes could produce fewer spikes at downstream targets (Khaliq and Raman, 2005; Monsivais et al., 2005). The effective output of an A1 cell would then depend not only on its somatic firing rate but also on the temporal pattern of its activity (Lisman, 1997; Bucher and Goaillard, 2011). Such activity-dependent propagation could filter the spike train between its initiation and its influence on distant retinal circuits, altering the timing and number of signals delivered to downstream neurons (Debanne et al., 1997, 2011). Future studies will be needed to determine whether these changes reflect propagation failure, uniform amplitude scaling, or a combination of mechanisms, and to establish their implications for signal transmission along A1 axonal arbors (Debanne et al., 1997, 2011).

## ACKNOWLEDGMENTS

We thank S. Cunnington for technical assistance. Tissue was provided by the Tissue Distribution Program at the Washington National Primate Research Center (WaNPRC; supported through NIH grant P51 OD-010425) and we thank the WaNPRC staff, particularly C. English and A. Baldessari, for making these experiments possible. This work was supported in part by grants from the NIH (NEI R01-EY027323 to M.B.M.; NEI R01-EY029247 to E.J. Chichilnisky, F.R., and M.B.M.; NEI R01-EY028542 to F.R.; R01-EY036594 to F.R., G. Field, J. Zylberberg, M.B.M.; NEI P30-EY001730 to the Vision Core) and the Research to Prevent Blindness Unrestricted Grant (to the University of Washington Department of Ophthalmology).

## AUTHOR CONTRIBUTIONS

S.A.B.: investigation, data curation, formal analysis, writing – original draft preparation, and visualization. F.R.: methodology, investigation, data curation, writing – reviewing and editing, resources, supervision, project administration, and funding acquisition. M.B.M.: conceptualization, methodology, investigation, software, formal analysis, data curation, writing – reviewing and editing, resources, visualization, supervision, project administration, and funding acquisition.

## METHODS

### Tissue preparation

Experiments were performed using an in vitro, pigment-epithelium attached preparation of the macaque monkey retina from three different macaque species of either sex (Macaca fascicularis, mulatta, and nemestrina) (Puller et al., 2015; Manookin et al., 2018). Tissues were obtained from terminally anesthetized animals that were made available through the Tissue Distribution Program of the National Primate Research Center at the University of Washington. All procedures were approved by the University of Washington Institutional Animal Care and Use Committee. Recorded cells were located in mid-peripheral or peripheral retina (4–8 mm, 18–36 degrees foveal eccentricity) with conditions closely matched to MEA recordings.

### Patch-clamp recording

Pieces of pigmented epithelium-attached retina were mounted ganglion cell-side up onto a poly-D-lysine-coated coverslip (BD Biosciences) and placed in a dish perfused with warmed (~32°C–34°C) bicarbonate Ames solution at ~6–8 mL min^−1^. Data were acquired using a Multiclamp 700B amplifier (Molecular Devices), digitized using an ITC-18 analog-digital board

(HEKA Instruments), and saved to disk using the Symphony data acquisition software (http://symphony-das.github.io).

### Multi-electrode array recording

Multi-electrode data was collected using custom arrays containing 512 electrodes. Electrodes were spaced at a 60 µm pitch in a 16 × 32 triangular arrangement, covering a rectangular region of approximately 1 × 2 mm, corresponding to approximately 4 × 8° of visual angle (Litke et al., 2004). The retina was placed ganglion-cell side down on the array and secured with a permeable membrane attached to a titanium plug. During recording, the retina was continuously perfused with oxygenated Ames’ solution maintained at 31–33°C. Voltage signals were band-pass filtered, amplified, and digitized at 20 kHz using custom electronics. Spikes were sorted using Kilosort (Pachitariu et al., 2024). Each unit was then evaluated using three quality-control criteria. First, units were excluded for evidence of contamination if they exhibited refractory-period violations within 1.5 ms or a receptive field containing a mixture of the receptive fields of other cells. Second, units with similar electrical images were treated as duplicate detections and merged. Third, units were cross-identified across stimulus sequences by correlating their electrical images.

### Visual stimulation

Visual stimuli were generated using the Symphony (http://symphony-das.github.io) and Stage (http://stage-vss.github.io) software packages and displayed on either an OLED monitor (eMagin) or a customized digital light projector (Lightcrafter 4500, Texas Instruments) (Appleby and Manookin, 2020, 2019). Stimuli were presented at medium to high photopic light levels with average L/M-cone photoisomerization rates (R*) of 1.5 × 10^4^ – 5.0 × 10^5^ s^−1^. The stimulus generation code for this manuscript is available on GitHub (https://github.com/mikemanookin/manookin-package).

### Analysis

#### Tissue, injections and immunohistochemistry

To recover cellular morphology following recording, we included 0.5–3.0% biocytin hydrazide (ThermoFisher) or neurobiotin (Vector Labs) in the internal recording solution to allow post hoc morphological recovery following fixation (4% paraformaldehyde) and immunostaining (streptavidin, Alexa Fluor 488; ThermoFisher). Cellular nuclei (DAPI) and starburst amacrine cell dendrites (anti-ChAT) were also labeled to localize the A1 dendrites within the inner plexiform layer (Manookin et al., 2008). Ankyrin G immunoreactivity was used to identify axon initial segments. The following antibodies were used: mouse anti-ankyrin G (1:500, Santa Cruz Bio., sc-12719), goat anti-ChAT (1:200, Millipore, AB144P), and DAPI (1:1000, ThermoFisher). Secondary antibodies included donkey anti-mouse Alexa Fluor 568 (1:500, ThermoFisher) and donkey anti-goat Alexa Fluor 568/633 (1:500, ThermoFisher). Anatomical verification was performed by taking images of filled cells using a Leica SP8 confocal microscope.

### Receptive fields from spatiotemporal noise

Receptive fields were measured with binary spatiotemporal noise consisting of a grid of 90–150 µm pixels jittered on a 20-30 µm grid. On each frame, every pixel took a contrast of +1 or −1 with equal probability, drawn from a pseudorandom sequence whose seed was stored with each epoch, so the stimulus could be regenerated exactly for subsequent analysis. We computed the spike-triggered average (STA) as the cross-correlation between the spike train and the contrast time course of each pixel and color channel. Spikes were counted in bins aligned to the frame times recorded with each epoch, two bins per frame, and the stimulus was regenerated from its seed at the same resolution. For pixel *i* with contrast *s_i_*(*t*) and spike count *r*(*t*), the cross-correlation at lag *τ* is *C_i_*(*τ*) = Σ*_t_ r*(*t*) *s_i_*(*t* − *τ*), the sum of the stimulus values that preceded each spike by *τ*. We evaluated it for *τ* = 0 to 60 bins (0–500 ms) and summed it across epochs.

For single-cell recordings the sum was computed in the Fourier domain (Baccus and Meister, 2002), and for array recordings as a matrix product over all cells of a recording at once (PyTorch); the two computations are mathematically equivalent. Because pixel contrasts are independent across pixels and frames and have zero mean, the STA estimates the cell’s linear filter up to a scale factor, at the spatial and temporal resolution of the stimulus (Chichilnisky, 2001). Only the shape of the STA is used here, and figures show it normalized to its largest absolute value. Each cell’s spatiotemporal receptive field was factorized by singular value decomposition into a spatial map and a temporal kernel. We took the contour at 0.30 of the peak of the spatial map as the receptive-field outline.

### Cross-correlograms

We computed cross-correlograms between the spike trains of each cell pair in 1-ms bins. Part of the correlation between two cells reflects their shared visual input. To separate correlations arising from shared visual input from those arising from interactions between cells, spatiotemporal noise was presented in consecutive pairs of epochs that used the same seed. A new random seed was drawn every other epoch, so each seed was presented exactly twice, in back-to-back epochs. The two epochs of a pair therefore delivered an identical stimulus, whereas trial-to-trial variability in the responses was independent between them. For each pair of cells, we computed the raw cross-correlogram from spikes recorded in the same epoch. We computed a stimulus-driven correlogram by pairing one cell’s spikes in the first epoch of a repeat pair with the other cell’s spikes in the second epoch, and vice versa, and averaging the two. Because the two epochs share the stimulus but not the trial-to-trial variability, this correlogram contains only the correlation produced by the common stimulus. Subtracting it from the raw correlogram leaves the correlation due to shared noise and direct interactions such as electrical coupling (Brivanlou et al., 1998; Trong and Rieke, 2008).

### Cell-type assignment on the array

We assigned array units to types from their receptive-field mosaic, electrical image, inter-spike interval distributions, and response kinetics, then matched them to patch-clamp recordings of anatomically identified cells. An electrical image with a single propagating track identifies a ganglion cell; several radiating tracks identify a polyaxonal amacrine cell (Greschner et al., 2014; Li et al., 2015).

### Static nonlinearities

Responses were described with a linear–nonlinear (LN) model, in which the stimulus is first filtered by the cell’s spatiotemporal receptive field and the result—the linear prediction or generator signal—is then passed through a static nonlinearity that converts it into a firing rate (Chichilnisky, 2001). The linear prediction at each stimulus frame *t* was the convolution of the stimulus with the receptive field, *g*(*t*) = Σ*_x_*_,*y*,*τ*_ *k*(*x*,*y*,*τ*) *s*(*x*,*y*,*t* − *τ*), where *k* is the spike-triggered average and *s* the stimulus contrast relative to the mean background. The nonlinearity was estimated without assuming a functional form: frames were grouped into bins by their linear prediction, and for each bin we computed the mean linear prediction and the mean spike rate in the same frames. The nonlinearity is the mean rate plotted against the mean prediction across bins. Because the receptive field was normalized, the prediction’s scale is arbitrary; the shape of the nonlinearity, and in particular whether the rate rises at negative as well as positive predictions, is what we compared across cells.

### Sparse noise

The sparse noise stimulus consisted of a grid of 90-150 µm pixels jittered on a 30 µm grid. On each frame, 1% of pixels were selected at random and set to a contrast of +1 or −1, with equal probability. All other pixels remained in the background. Each frame was presented for 200 ms. Spike counts were binned in register with the frame updates, and the receptive field was estimated as the spike-triggered average of the stimulus frames. For each frame, filtered contrast was the projection (inner-product) of the frame onto this receptive field. Because spikes were binned frame by frame, responses evoked when a pixel returned to the background fell in the following frame, where filtered contrast was typically near zero.

*Nonlinearity calculation.* The linear prediction was computed with the spike-triggered average estimated from the same sparse-noise recording. Stimulus frames and spike counts were averaged within each 200-ms frame (12 display frames at 60 Hz), so each sample paired one stimulus frame with the spikes it evoked. The prediction was divided by its standard deviation, the frames were sorted by prediction and divided into 25 bins containing equal numbers of frames, and the mean spike count in each bin was converted to a rate (spikes s^−1^).

### Natural movies

Natural movies from the DOVES database (Van Der Linde et al., 2009) were presented at 60 Hz for 6 seconds on each trial. Linear predictions were computed by convolving each movie with the spatiotemporal receptive field measured with white noise (see Figure 1). Nonlinearities were then estimated as above.

*Nonlinearity calculation.* The receptive field was the spike-triggered average from the spatio-temporal noise (see Figure 1), with 61 lags (1 s) at 60 Hz. Its DC offset was removed by subtracting the mean of the last third of the lags, pixels whose peak amplitude was below 3 robust standard deviations of the noise were set to zero, and the kernel was scaled to unit norm. Each movie was reconstructed on the receptive-field grid and convolved with the full spatiotemporal kernel. Spikes were counted in bins matching the 60-Hz display frames and averaged across repeats of the same movie. The first part of each movie (the initial fixation period plus the length of the filter, during which the convolution is incomplete) was discarded, and all movies a cell saw were pooled. The linear prediction was divided into 9 bins of equal width spanning approximately the 1st to 99th percentile, and bins containing fewer than 15 frames were excluded. Equal-width rather than equal-count bins were used here because the two arms of an On–Off nonlinearity lie at large |prediction| values, where equal-count bins are widest and average the tails into the center of the distribution.

### Expanding-spot stimulus and area summation

We presented dark spots of 18 diameters from 18 to 912 µm, centered on the receptive field, at ±0.5 contrast for 0.5 seconds against a mean background, and in most cells bright spots of the same diameters. We averaged the spike rate over a 200-ms window after spot onset and fitted each resulting area-summation curve with a difference-of-Gaussians model (Rodieck, 1965; Enroth-Cugell and Robson, 1966).

### Receptive-field center diameter

We defined the center diameter as the spot diameter at which the curve first reaches 1 − 1/e of its maximum, as determined by the difference-of-Gaussians fit. The On curve was measured at the onset of bright spots and the Off curve was measured at the onset of dark spots. We compared types with a Kruskal–Wallis test and compared On against Off centers within cells with a two-sided Wilcoxon signed-rank test.

### Chirp stimulus

To characterize temporal response properties and contrast sensitivity in a single presentation, we used a “chirp” stimulus modeled on Baden et al. (2016). The stimulus was presented on a photopic mean background. Each trial consisted of a fixed sequence of five segments separated by 1 s at the background intensity: (1) a 500 ms pre-stimulus period at background; (2) a 500 ms light increment and, after a 1 s gap, a 500 ms decrement, each at 100% Weber contrast; (3) a 10 s sinusoidal frequency sweep at 100% contrast in which the temporal frequency increased linearly from 0 to 10 Hz; and (4) an 8 s sinusoidal contrast sweep at a fixed 2 Hz carrier in which contrast increased linearly from 2% to 100%. The trial ended with a 500 ms tail period. Total trial duration was 22.5 s.

### Analysis of chirp responses

Cellular responses were separated according to the known segment boundaries of the stimulus. For the steps we measured the mean rate in the 200 ms after the onset of each step and subtracted the mean firing rate prior to stimulus onset. The On-Off index (Carcieri et al., 2003) is (I − D)/(I + D), where I is the response to the light-step onset and D the response to the dark-step onset. We used only the onsets because both start from the mean background, whereas the responses at the step offsets follow adaptation to the preceding step. For the frequency sweep, the response was divided into individual stimulus cycles using the zero crossings of the regenerated stimulus waveform; for each cycle we computed the instantaneous stimulus frequency and the response amplitude as the fundamental (F1) Fourier component. F1 amplitudes were averaged in 1-Hz bins, and temporal tuning was summarized by the ratio of the mean F1 at 8–10 Hz to that at 1–3 Hz, both within the 0–10 Hz range swept in every recording. For the contrast sweep, cycle-by-cycle response amplitudes at the 2 Hz carrier were plotted against instantaneous contrast. The half-maximal contrast (C_50_) was measured without a model: the curve was made monotonic by isotonic regression, and C_50_ is the contrast at which it first reached half its maximum (linear interpolation), so it always lies within the tested range. Population data are reported as mean ± SEM across cells. Each summary measure was compared across the four types with a Kruskal–Wallis test, and each type with Off parasol cells with a two-sided rank-sum test. Maintained firing rates (rate at a mean photopic background) of A1, broad thorny and recursive bistratified cells were pooled and compared with those of Off parasol cells with a two-sided rank-sum test.

### Motion stimuli

We presented drifting gratings and moving bars at 8 to 12 directions spanning 360°, in randomized order. Gratings were presented across a range of spatial (0.25–4 cycles degree^−1^; half-cycle width, 50–800 μm) and temporal frequencies (0.5–8 Hz; contrast, 0.5–1.0). Bars were moved in 12 different directions across a range of speeds (600–1200 μm s^−1^; bar width, 300–600 μm).

### Selectivity indices

We computed the direction-selectivity index as the length of the vector sum of the per-direction mean firing rates, divided by their sum. The orientation-selectivity index uses the same construction on doubled angles. Both are bounded between 0 and 1, and both are biased upward by noise when firing rates are low or repeats are few (Mazurek et al., 2014). We compared types with a Kruskal–Wallis test. Statistics on the array data use the bias-corrected index: for each cell we subtracted the mean of its own shuffled null, obtained by permuting the per-direction responses. The correction is needed because the raw index is biased upward when firing rates or trial counts are low, and that bias differs systematically across types. Off parasol cells fire at the highest rates and make up by far the largest sample, so with the raw index they sit closest to the true floor, and types that fire less appear more selective than they are.

### Burst detection

We detected bursts as isolated triplets. According to our formulation, a burst began when the two intervals after it were each shorter than 30 ms and the interval before it was at least 30 ms. That spike and the two following it became the first, second and third spikes of the burst. The isolation requirement makes the triplets non-overlapping, so no spike is counted both as a third spike and as a first. Where a high-frequency run held more than three spikes, we used only the first three, so that the first-to-third comparison always spans the same ordinal positions. A burst therefore lasts at most 60 ms.

### Electrical images and simulated propagation failure

For each cell we computed electrical images separately from the first spikes of bursts, the third spikes of bursts, and all spikes. We zero-centered each spike’s array snapshot by subtracting the mean signal over a 3-ms segment ending 4 ms before that spike, which removes slow drift without touching the spike. We compared images by Pearson correlation across all electrodes and samples. To ensure a sufficient signal to noise ratio for burst electrical images, only cells with a minimum of 3000 burst spikes were included in the analysis.

We simulated a propagation failure at the level of a single spike, replacing that spike’s array data with the data recorded 4 ms earlier. This removes the spike’s contribution to the average and preserves the recording’s noise. We added failures one spike at a time, in random order, and recomputed the correlation between the first-spike image and the full image after each. The reported count is the number of failures at which that correlation falls to the value already shown by the third burst spike, averaged over 20 simulations per cell. Expressing it as a fraction of the cell’s spikes makes it comparable across cells with different firing rates.

### Statistics

Population values are reported as mean ± SEM unless stated otherwise, and *n* is the number of cells. Comparisons across cell types used the Kruskal–Wallis test, pairwise comparisons the two-sided Wilcoxon rank-sum (Mann–Whitney U) test, and within-cell comparisons the Wilcoxon signed-rank test; tests are two-sided and unadjusted unless noted, and where several comparisons are made within one figure the conservative Bonferroni threshold is stated. Permutation tests are described with the figure that uses them. No statistical test was used to determine sample size.

